# Dissociable neural correlates of multisensory coherence and selective attention

**DOI:** 10.1101/2022.02.01.478616

**Authors:** Fei Peng, Jennifer K. Bizley, Jan W. Schnupp, Ryszard Auksztulewicz

## Abstract

Previous work has demonstrated that performance in an auditory selective attention task can be enhanced or impaired, depending on whether a task-irrelevant visual stimulus is temporally coherent with a target auditory stream or with a competing distractor. However, it remains unclear how audiovisual (AV) temporal coherence and auditory selective attention interact at the neurophysiological level. Here, we measured neural activity using electroencephalography (EEG) while participants performed an auditory selective attention task, detecting deviants in a target audio stream. The amplitude envelope of the two competing auditory streams changed independently, while the radius of a visual disc was manipulated to control the audiovisual coherence. Analysis of the neural responses to the sound envelope demonstrated that auditory responses were enhanced independently of the attentional condition: both target and masker stream responses were enhanced when temporally coherent with the visual stimulus. In contrast, attention enhanced the event-related response (ERP) evoked by the transient deviants, independently of AV coherence. Finally, we identified a spatiotemporal component of the ERP, likely originating from the superior temporal gyrus and the frontoparietal network, in which both attention and coherence synergistically modulated ERP amplitude. These results provide evidence for dissociable neural signatures of bottom-up (coherence) and top-down (attention) effects in the AV object formation.

## Introduction

In many real world sound environments, sounds originate from multiple sources – the auditory system needs to appropriately segregate and group sound features to efficiently process the entire scene (Maddox & Shinn-Cunningham, 2012; Middlebrooks et al., 2017; Shamma et al., 2011). Several psychoacoustic studies have demonstrated that visual cues which are temporally coherent with sounds can facilitate auditory processing. For example, a synchronous, task-irrelevant light flash improves the detection of weak auditory signals (Lovelace et al., 2003). Similarly, task-irrelevant visual stimuli which are temporally coherent with a speech envelope enhance speech intelligibility in background babble noise (Yuan et al., 2020). Furthermore, performance in an auditory selective attention task can be enhanced or impaired, depending on whether the task-irrelevant visual stimulus is temporally coherent with a target sound stream or a competing masker stream (Atilgan & Bizley, 2021; Maddox et al., 2015). These findings suggest that temporal coherence between auditory and visual stimuli can facilitate binding of audiovisual (AV) features to enable AV object formation (Bizley et al., 2016), and that the AV object might be easily captured by attention. However, the neural mechanisms mediating the interactions between temporal coherence and selective attention in facilitating AV integration remain unknown.

Several previous studies have identified potential neural correlates of attentional modulation of AV integration. For example, a study using simple tone pips and visual gratings demonstrated that ERPs related to multisensory integration were amplified by selective attention (Talsma & Woldorff, 2005). A follow-up study showed that when both visual and auditory stimuli were attended, the ERP peak amplitude showed superadditive AV effects; however, subadditive effects were observed for unattended stimuli (Talsma et al., 2007). In order to go beyond relatively simple, discrete stimuli, some EEG and MEG studies have employed the analysis of “neural envelope-tracking responses” to speech, by modeling the relationship between neural activity and the auditory envelope it represents (Crosse et al., 2015; Golumbic et al., 2013), and have found that congruent audio-visual speech enhances the envelope tracking response relative to auditory speech alone or the linear summation of auditory and visual speech. Other studies have used auditory selective attention tasks to show that attention is necessary for AV speech integration. For example, Morís Fernández et al. (2015) recorded fMRI data while two competing continuous speech streams were presented simultaneously with a video speech stream, and participants were asked to attend or ignore the speech congruent with the video. Although the physical stimuli in the two conditions were identical, multisensory integration was found to occur almost exclusively only when the congruent AV speech was attended. However, another study by Ahmed et al., (2021) found some evidence for early AV integration in the unattended stream, consistent with the idea that distinct audiovisual computations emerge at different processing stages (Kayser & Shams, 2015; Talsma et al., 2007; Talsma & Woldorff, 2005; Zumer et al., 2020). One potential difficulty with interpreting findings from AV speech processing is that it can be hard to know the extent to which they generalize to other continuous AV stimuli, given that speech processing can be heavily influenced by linguistic knowledge and expectations. The McGurk effect provides a well-known example, in which a visual syllable /ga/ presented synchronously with an auditory /ba/ can produce fusion percepts such as /da/ or /tha/, which reflect neither of the conflicting visual and auditory inputs faithfully, but rather correspond to a phonetically plausible compromise (McGurk & McDonald 1976). These kinds of fusion percepts which combine multisensory processing with implicit phonetic knowledge are very interesting, but they may not be representative of more general mechanisms of visual influences on auditory processing.

Bizley et al., (2016) proposed that AV binding is a form of AV integration that underlies perceptual object formation. A recent neurophysiological study in ferrets (Atilgan et al., 2018) provided direct evidence that temporal coherence between features of audio and visual stimuli facilitates cross-modal binding in single neurons in the auditory cortex. The study found that AV temporal coherence not only enhanced the neural representation of the binding features (i.e., the coherent auditory amplitude and visual luminance changes), but also of the other (non-binding) sound features, including timbral changes that were not predicted by the visual stimulus. It remains unclear whether such bottom-up effects modulate the cortical representation of auditory streams independently of attentional top-down enhancement of sound stream processing, or whether, and where in the brain, the two effects interact.

Here, we use EEG to investigate the electrophysiological correlates of the effects of AV temporal coherence and auditory selective attention on sound processing. Our aims were two-fold: first, to investigate how AV coherence and attention affected the neural signatures of continuous stream processing as manifest in the envelope-tracking response, and second, to examine their effects on the neural responses to the transient auditory deviants that subjects were cued to detect or ignore. We hypothesized that the effects of temporal coherence and attention might be independent but synergistic.

## Results

During the EEG recording, participants performed an auditory selective attention task adapted from previous studies (Atilgan & Bizley, 2021; Maddox et al., 2015). In each trial the auditory stimuli consisted of two streams: an attended stream, which will be referred to as the auditory target (At), and a to-be-ignored stream, which will be referred to as the auditory masker (Am). Each stream was constructed by amplitude-modulating a continuous artificial vowel stimulus, either a /u/ sound with a lower pitch of 175 Hz, or an /a/ sound with a slightly higher pitch of 195 Hz (the target stream identity was randomised and counterbalanced). On each trial, the target stream started 1 s earlier than the masker, which cued to the participants which stream to attend to during that trial. In addition to the two auditory streams, participants were presented with a concurrent visual stream (V) consisting of a radius-modulated disc (that is, a filled circle that continually contracted and expanded). The two auditory streams were independently amplitude-modulated, and the modulation of the visual disc could be temporally coherent with either the amplitude of the target stream (AtAmVt), or the masker stream (AtAmVm), or could vary independently of both (AtAmVi) (Figure 1A). At random time points, auditory “timbre deviants” were introduced in both target and masker streams. During these 200 ms deviants, the /u/ briefly changed to an /ε/ and back, or the /a/ changed to an /i/. Participants were asked to detect auditory deviants embedded in the target auditory stream by pressing a keyboard button, while ignoring the masker stream deviants. Each trial lasted 14 s, and the stimuli were presented in 12 blocks of 18 trials. A ‘hit’ was defined as the response to the deviant in the target auditory stream within 1 s following the onset of the deviant, and a ‘false alarm’ was defined as the response to a deviant that occurred in the masker stream.

**Figure 1.**
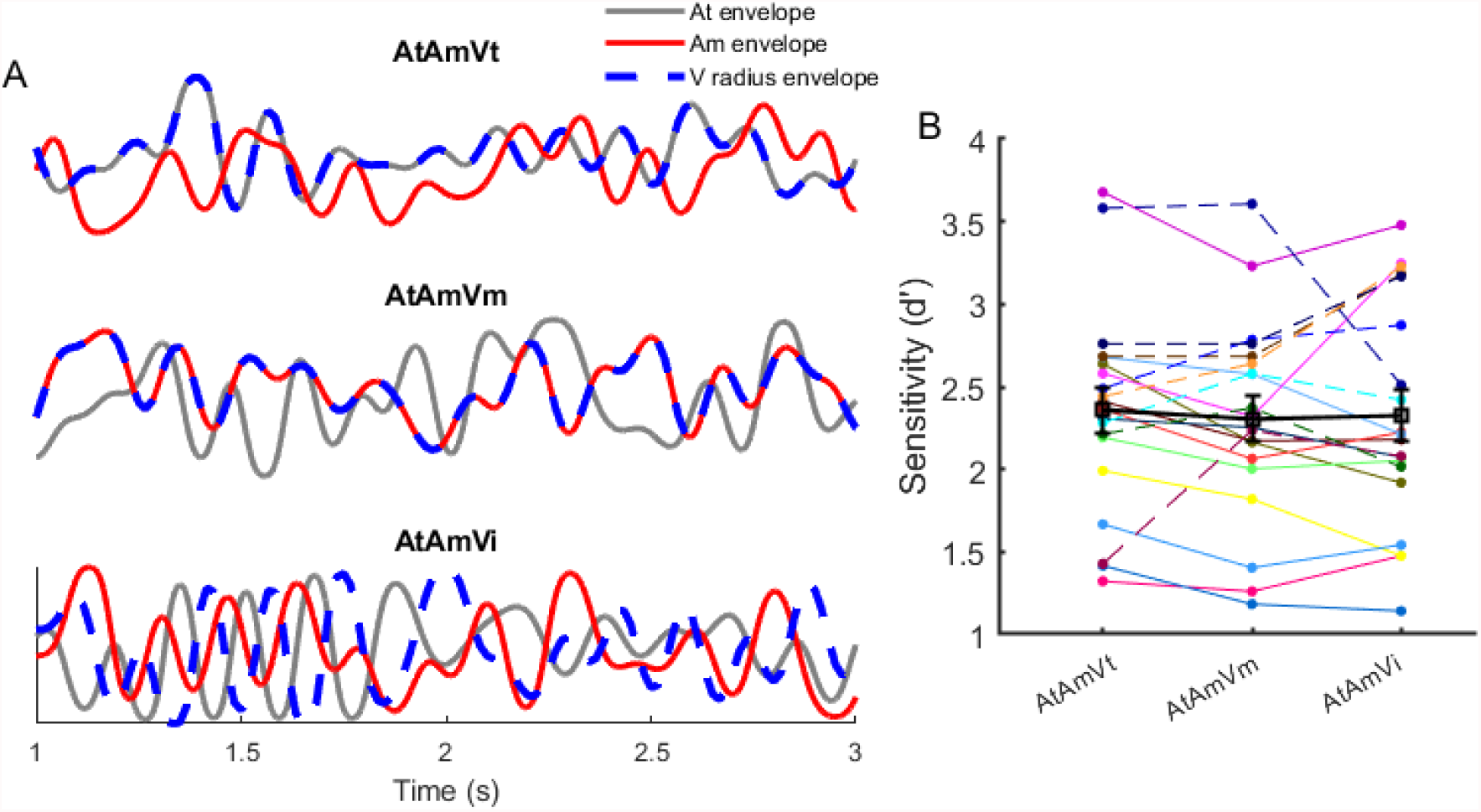
Experimental paradigm and behavioral performance. **(A)** Schematic plot of auditory and visual stimuli in the behavioral task. Amplitude envelopes of target/attended sound (grey solid line), masker/unattended sound (red solid line), and visual radius envelope (blue dashed line). **(B)** Behavioral sensitivity (d’) for each visual condition. Each line shows data of one participant. Solid lines indicate participants with higher d’ in the AtAmVt vs. AtAmVm condition, and dashed lines indicate participants with lower d’ in the AtAmVt vs. AtAmVm condition. Black squares represent group averages, and error bars indicate the standard error of the mean (SEM).

### Behavioral results

First, we investigated whether behavioral performance was stable over time, which would warrant pooling data from all blocks. To this end, we calculated the single-participant hit rate separately for each of the 12 blocks, and fitted the data using a linear regressor representing the block number. The resulting regression coefficient (slope) was not statistically different from zero across participants (one sample t-test, p = 0.28), suggesting that there were neither significant learning nor fatigue effects during the experiment.

To investigate the effect of the visual temporal coherence on behavioral performance, we performed one-way repeated measures ANOVAs on d’ (Figure 1B), hit rates and false alarm rates. No significant effect of visual coherence on deviant detection was observed, likely due to large variability and heterogeneity of response patterns across participants. For instance, while some participants showed behavioral benefits of visual coherence (e.g., larger d’ in AtAmVt condition than AtAmVm), others showed the opposite effects (Figure 1B). Two previous studies using similar stimulus paradigms (Atilgan & Bizley, 2021; Maddox et al., 2015) reported enhanced task performance when the target stream and visual stimulus were temporally coherent. Our failure to replicate these data may be attributable to small but perhaps important differences in the details of the experimental paradigms (see Discussion). However, the aims of this study were to identify effects of attention and AV coherence on physiological measures of neural stimulus representations, and the timbre deviants primarily served as a device for controlling and monitoring our participants’ attention. The relatively high hit rates and low false alarm rates indicate that the deviants had fulfilled that purpose.

### Stimulus reconstruction reveals temporal coherence mediated audiovisual integration

To investigate the occurrence of AV integration at both attended and unattended conditions, we reconstructed an estimation of the sound envelope from the recorded EEG waveforms. We used the condition in which the visual stimulus was independent of both auditory streams to estimate unimodal reconstructions for the target auditory stream (At), the masker stream (Am) and the visual stream (Vi) (Figure 2B). The unimodal reconstructions for all conditions were significantly better than chance derived by the permutation test. We found that the average reconstruction accuracy of Vi was significantly higher than At (Wilcoxon signed-rank test p = 0.002) and Am (Wilcoxon signed-rank test p = 0.001). From these we estimated the response to stimuli in which the visual stimulus was coherent with one or the other stream by linear summation. This linear summation model was compared to an integration model in which audiovisual envelopes were reconstructed based on the responses to conditions in which the visual stimulus was temporally coherent with one or the other stream (i.e., AtVt and AmVm) (Figure 2A). We found that the average reconstruction accuracy of the AV decoder was significantly higher than that of the A+V decoder for both the target stream (Figure 2C, Wilcoxon signed-rank test p < 0.001) and the masker stream (Figure 2D, Wilcoxon signed-rank test p < 0.001), consistent with AV integration occurring. However, no significant difference was found between the reconstruction accuracy of the target and masker sound using the AV decoder, suggesting that while visual information dominated the stimulus reconstruction AV integration was occuring independently of attention (Wilcoxon signed-rank test p = 0.455). To further investigate how the visual coherence and attention affect the reconstruction at individual time-lags, the single-lag reconstruction was also compared between the AV decoder and the A+V decoder. The reconstruction accuracy of the AV decoder was significantly larger than the A+V decoder across all the time lags for both target (Figure 2E) and masker streams (Figure 2F) (cluster level p_FWE_ <0.001), supporting the conclusion that temporal coherence facilitates AV integration independently of attention.

**Figure 2.**
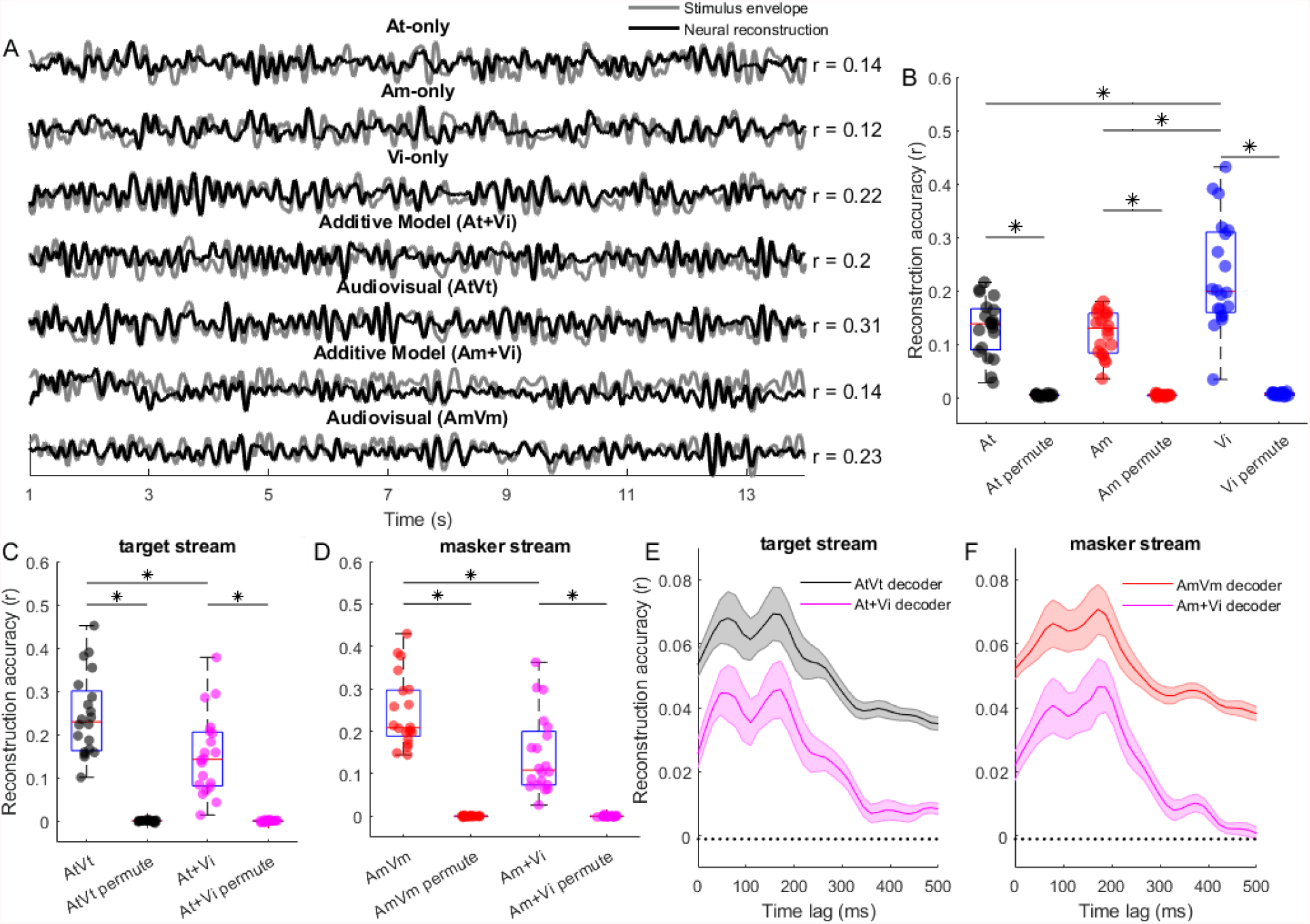
Stimulus reconstruction. **(A)** Examples of the original sound envelope (grey) with the grand-average neural reconstruction (black) overlapped. The mean reconstruction accuracy over subjects is indicated to the right. **(B)** The stimulus reconstruction accuracy for each stream in the independent condition AtAmVi was significantly better than chance (permutation test). Each dot represents one subject. **(C, D)** The stimulus reconstruction accuracy using the AV decoder and A+V decoder for the target and masker sound was significantly better than chance (permutation test), respectively. **(E, F)** The single-lag reconstruction accuracy using the AV decoder and A+V decoder for the target and masker sound, respectively.

### Forward models highlight attentional modulation of auditory responses

We next asked how temporal coherence and attention affect AV integration across the different EEG channels by estimating the temporal response functions (TRFs) of each channel. While stimulus reconstruction predicts the accuracy of cortical tracking of the amplitude envelope by using multichannel EEG response, TRFs reflect the brain’s linear transformation of the sound envelope to the neural responses at each EEG channel. We first explored whether we could observe similar evidence of audiovisual integration from the TRF estimations as we did with the stimulus reconstruction. We estimated unisensory TRFs for the auditory target stream (TRF_At_), the auditory masker stream (TRF_Am_), and the visual stimulus (TRF_Vi_), separately, from the response in the condition AtAmVi, in which temporal envelopes of all three streams were independent. We then estimated the TRF_AtVt_ and TRF_AmVm_ using the responses in the condition AtAmVt and AtAmVm, respectively.

We compared the TRF_AV_ amplitude with the linear summation of TRF_A_ and TRF_V_ amplitude over all channels and time lags using a paired t-test highlight AV integration, which would manifest as deviations from linear summation. When the visual stimulus was temporally coherent with the target stream (i.e., AtAmVt), there was no significant difference between the amplitude of the estimated TRF_AtVt_ and the linear summation of TRF_At_ + TRF_Vi_ across all the channels and time lags (Figure 3A). However, when the visual stimulus was temporally coherent with the masker sound (Am) condition, the analysis indicated that TRF_AmVm_ amplitude was significantly stronger than TRF_Am_ + TRF_Vi_ amplitude. This effect was observed over the central and frontal spatiotemporal channels between time lag 188 to 250 ms (Figure 3B, cluster-level p_FWE_ = 0.005, T_max_ = 3.71). For the masker stream, both the forward model (TRF estimation) and stimulus reconstruction yielded evidence for AV integration. However, for the target stream, only the stimulus reconstruction, but not the TRF, provided the evidence of AV integration.

**Figure 3.**
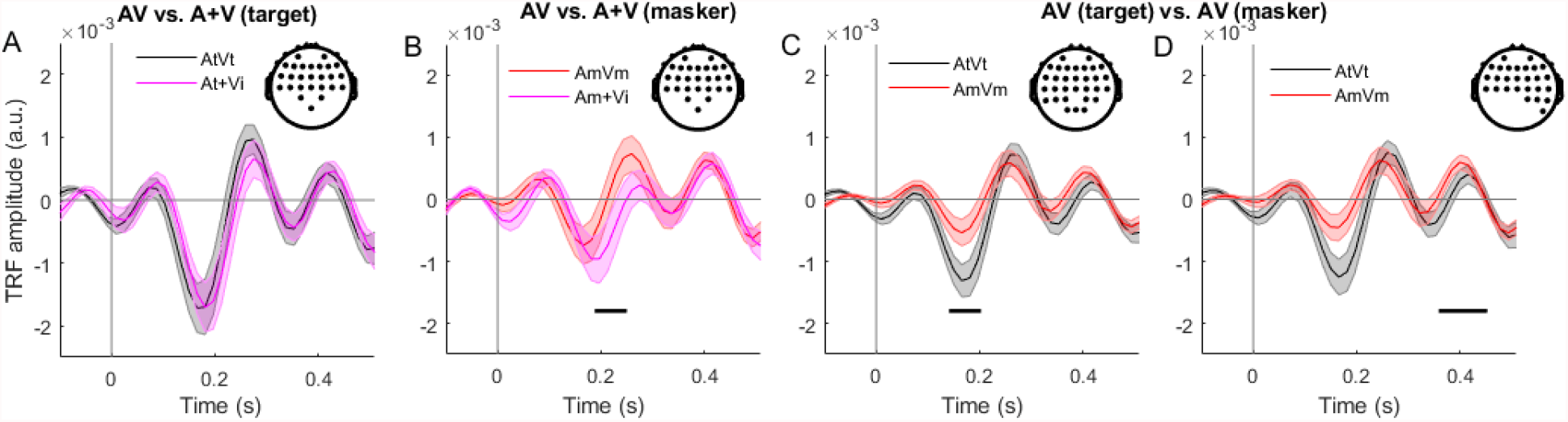
Temporal response function analysis. **(A)** For the target (At) sound condition, the TRF estimated for coherent AV streams was not significantly different from the summation of independent stream (At + Vi). **(B)** For the masker sound (Am) condition, the TRF estimated for coherent AV streams (AmVm) had a stronger amplitude than the summation of TRFs estimated for independent AV streams (Am + Vi). Shaded areas indicate SEM (standard error of the mean) across subjects. The topographical plot shows the EEG channel locations with a significant difference. Black horizontal bars: p_FWE_ < 0.05. **(C, D)** TRFs were modulated by attention for two clusters showing the significant difference at different time lags, distinguishing between AV streams in which the visual stream was coherent with the target sound (AtVt) vs. the masker sound (AmVm). Figure legend as in (A).

To investigate the effect of attention, independently of AV temporal coherence, on the cortical representation of AV amplitude envelopes, we compared the TRF_AtVt_ and TRF_AmVm_ amplitude using a paired t-test. Selective attention modulated the magnitude of the TRF: the TRF_AtVt_ amplitude was significantly larger than the TRF_AmVm_ amplitude (Figure 3C and Figure 3D). This effect was observed over two spatiotemporal electrode clusters: one over central and frontal channels between time lag 156 to 203 ms (Figure 3C, cluster level p_FWE_ < 0.001, T_max_ = 4.23), and one over frontal, and right temporal channels between time lag 359 to 391 ms (Figure 3D, cluster level p_FWE_ = 0.03, T_max_ = 4.79).

In summary, we observed evidence that AV integration occurred both in the target and masker auditory stream when measures of stimulus reconstruction accuracy were used to analyse the neural responses to the sound envelopes. Analysis of TRFs amplitude across all EEG channels showed that attention modulated the magnitude of the TRF. AV integration was observed for the masker stream in central and frontal channels. Taken together, our results suggest audio-visual integration occurs automatically, prior to attentional modulation.

### Effects of audiovisual temporal coherence and selective attention on deviant-evoked responses

The analysis so far has focused on the neural responses to the amplitude envelopes of the audiovisual scene, and has revealed evidence for both attentional modulation of acoustic responses, and AV integration of temporally coherent cross-modal sources. Since, in the temporally coherent conditions, the visual and auditory streams convey redundant information, this integration falls short of reaching the stricter definition of binding proposed by Bizley et al., (2016) which requires an enhancement of independent features that are not those which link the stimuli across modalities. Here, the presence or timing of the auditory timbre deviants that listeners detected in the selective attention task are not predicted by the amplitude changes of the audio or visual envelopes, and they thus provide a substrate with which to explore binding.

To investigate how AV temporal coherence and attention affect the deviant-evoked responses, we compared the ERPs evoked by deviants embedded in At and Am streams, and, in order to look for evidence of binding, asked how audiovisual temporal coherence modulated these responses (Figure 4 and Figure 5).

**Figure 4.**
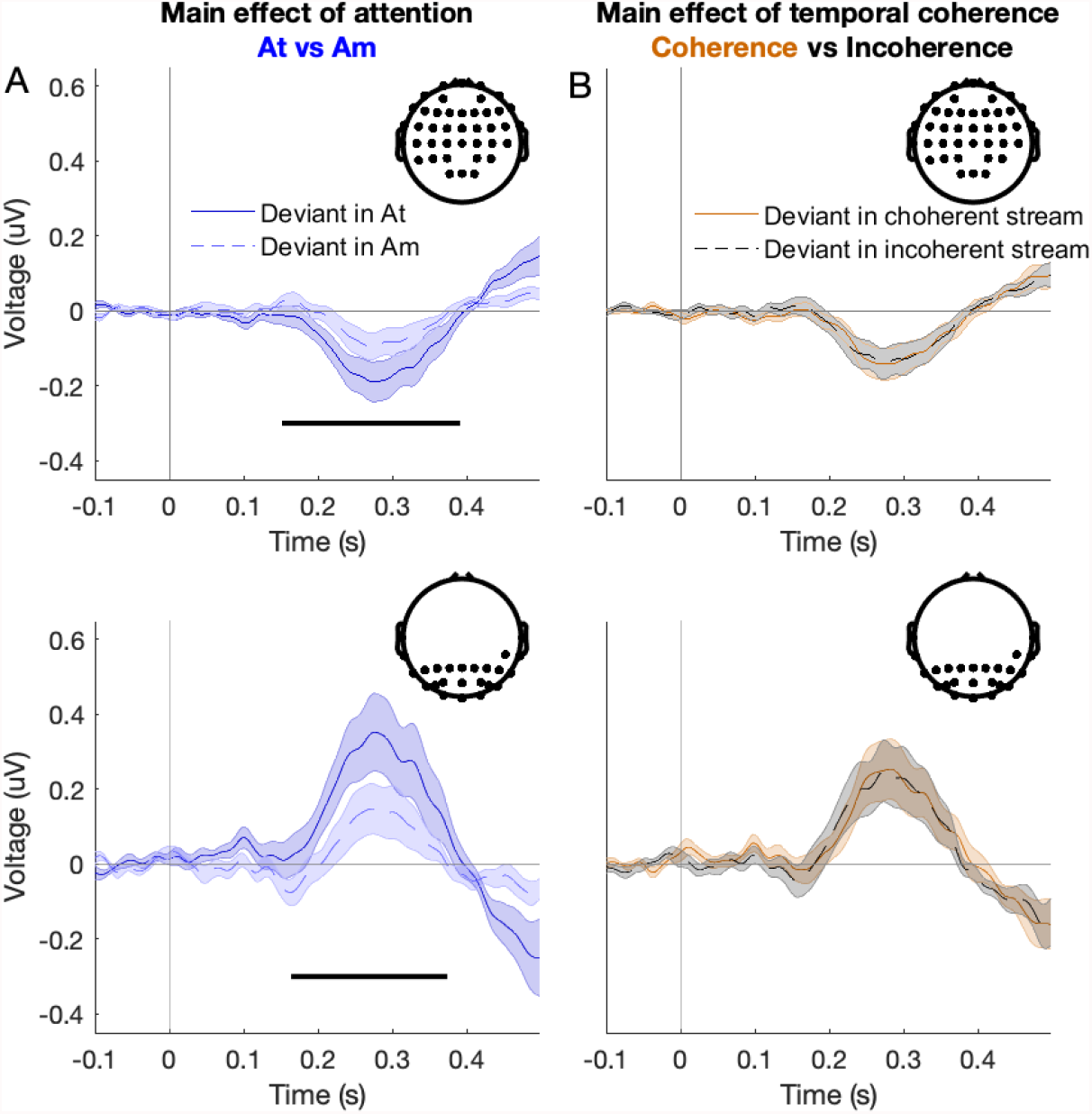
Attentional enhancement of deviant-evoked ERPs: channel-by-channel mass-univariate analysis. Traces show grand-average deviant-evoked ERPs in target (At) and masker (Am) streams, separately for each visual coherence condition: **(A)** Deviant-evoked ERPs in At (blue solid lines) and Am (blue dashed lines) averaged across coherence conditions. The topographical plot shows the EEG channel locations where the ERP amplitude difference between the two conditions (as indicated at the top of each plot) was significantly observed (FWE-corrected). The black bar represents the time segment with a significant difference between the deviants in two different conditions. **(B)** Deviant-evoked ERPs in coherent stream (yellow solid lines) and incoherent stream (black dashed lines) averaged across attention conditions. The topographical plot shows the EEG channels locations where the main effect of attention was significant (since no main effect of temporal coherence was observed). Shaded areas represent SEM across subjects.

**Figure 5.**
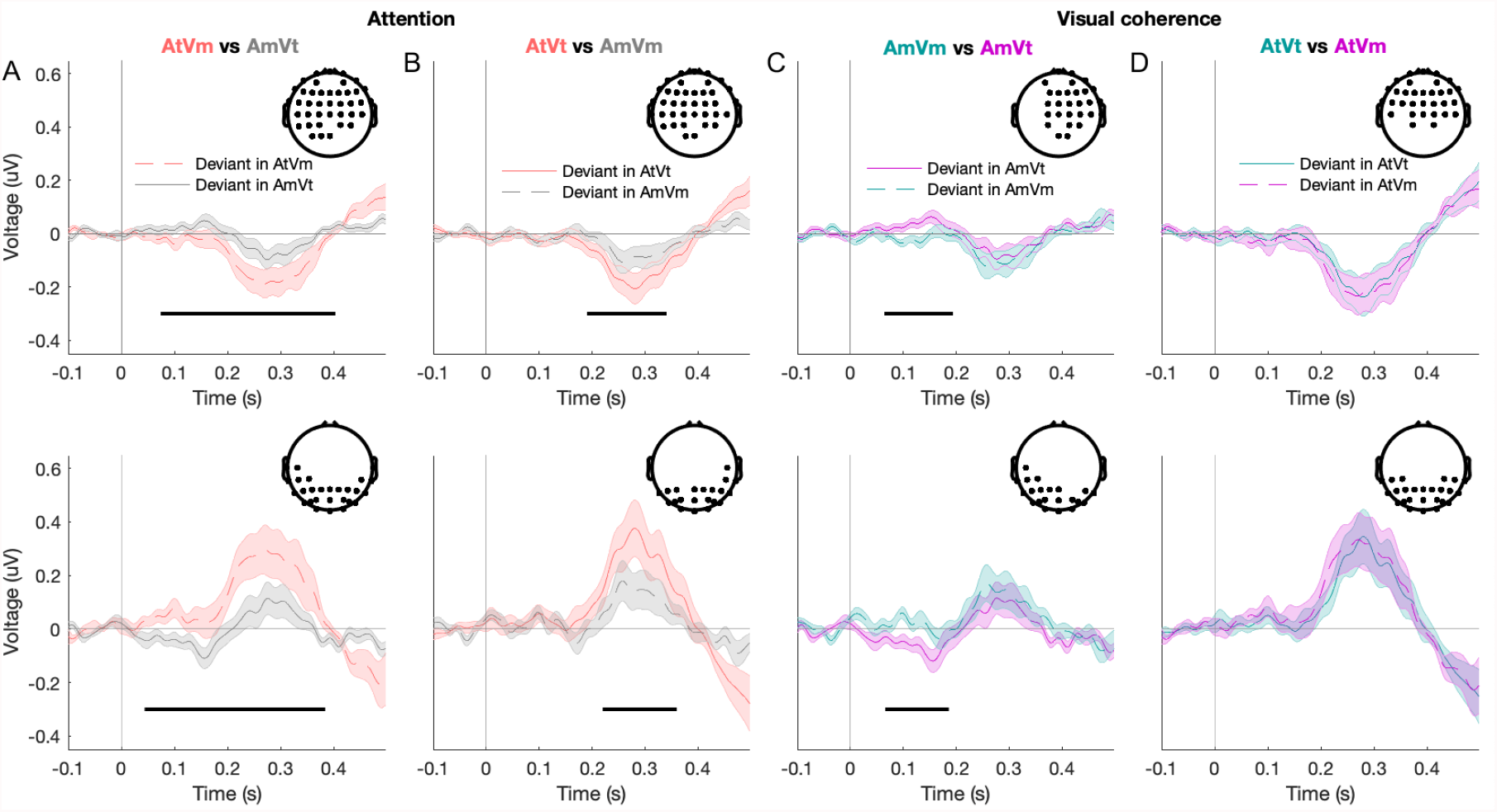
Grand-average deviant-evoked ERPs between condition comparisons. **(A)** Deviants presented in the incoherent target stream (red dashed lines) and masker stream (grey solid lines); **(B)** Deviants presented in the AV coherent target (red solid lines) and masker stream (black dashed lines); **(C**,**D)** Deviants presented in the masker and target stream in each of the two attentional conditions (left: masker stream; right: target stream); The topographical plots in panels (A-C) show the EEG channel locations where a significant ERP amplitude difference between the two conditions (as indicated at the top of each plot) was observed (FWE-corrected) except. The topographical plot in (D) shows the EEG channel locations where the interaction effect between attention and coherence was significant. The black bar represents the time segment with a significant difference between the deviants in two different conditions. Shaded areas represent SEM across subjects.

In a traditional channel-by-channel mass-univariate analysis, correcting for multiple comparisons across all channels and time points, we observed a significant main effect of attention (Figure 4A, anterior cluster, 248 to 378 ms, F_max_ = 16.43, Z_max_ = 3.61, p_FWE_ < 0.001; posterior cluster, 200 to 376 ms, F_max_ = 16.10, Z_max_ = 3.57, p_FWE_ < 0.001) and a significant interaction effect of attention and temporal coherence (anterior cluster, 210 to 254 ms, F_max_ = 14.88, Z_max_ = 3.43, p_FWE_ = 0.008). No main effect of temporal coherence was observed (Figure 4B).

Significant post-hoc comparisons between conditions were consistent with the main effect of attention: for both temporally coherent and temporally independent streams, the deviant response in the target always exceeded that of the masker. The amplitude of the ERP evoked by timbre deviants presented in the target stream (AtVm) was significantly larger than that in the masker stream (AmVt) in two clusters: negative peak enhancement was observed over anterior channels (Figure 5A the first row, 74-408 ms after deviant onset, p_FWE_ < 0.001, T_max_ = 4.23), and positive peak enhancement over posterior channels (Figure 5A the second row, 44-384 ms after deviant onset, p_FWE_ < 0.001, T_max_ = 4.68). In the AV coherent stream, we observed that ERP amplitude evoked by the timbre deviants in the attended coherent stream (AtVt) was significantly stronger than in the unattended coherent stream (AmVm) in two clusters: one over the central and frontal channels between time lag 190 to 348 ms (Figure 5B the first row, cluster level p_FWE_ < 0.001, T_max_ = 3.8), and one over posterior channels between time lag 220 to 360 ms (Figure 5B the second row, cluster level p_FWE_ = 0.007, T_max_ = 4.18).

Post-hoc comparisons also allowed us to examine the interaction between temporal coherence and attentional condition. We observed that the amplitude of ERP evoked by deviants in the masker stream was significantly smaller when this was accompanied by a temporally coherent visual stimulus (Figure 5C). The deviant induced ERP was smaller in the AmVm condition than in the AmVt condition in two clusters: one over the central and frontal channels between time lag 64 to 196 ms (Figure 5C the first row, cluster level p_FWE_ = 0.011, T_max_ = 4.38), and one over left temporal and posterior channels between time lag 66 to 190 ms (Figure 5C the second row, cluster level p_FWE_ = 0.005, T_max_ = 3.72). In contrast, audiovisual temporal coherence did not influence the size of the deviant response in the target stream (Figure 5D).

From the mass-univariate ERP data analysis, we did not observe any significant effect of the temporal coherence on the deviants evoked response. We hypothesized that the temporal coherence affects the spatiotemporal components of the neural responses. In a follow-up analysis, we investigated whether effects of attention and visual coherence can be mapped onto spatiotemporal components of variance in the ERP data. To this end, we performed a principal component analysis to extract the spatiotemporal component of the ERP, and performed separate two-way repeated measures of ANOVAs with two main factors: attention (attended and unattended) and visual coherence (coherent and incoherent), on the first four principal components (PCs) in the time domain. The analysis of the 1^st^ PC (Figure 6A, explaining 67% of the original variance) only showed a main effect of attention (time lag between 208 to 284 ms, F_max_ = 32.53, Z_max_ = 4.92, p_FWE_ < 0.001). No main or interaction effects were found to be significant for the 2^nd^ and 4^th^ PC (Figure 6B and Figure 6D, explaining 6% and 3% of the original variance, respectively). However, the analysis of the 3^rd^ PC (explaining 4% of the original variance) showed a main effect of attention (time lag between 8 to 170 ms, F_max_ = 43.33, Z_max_ = 6.88, p_FWE_ < 0.001), and the interaction effect between attention and visual coherence (time lag between 214 to 238 ms, F_max_ = 14.82, Z_max_ = 3.43, p_FWE_ < 0.001). We observed an effect of attention on the 3^rd^ PC amplitude only when the visual stimulus was coherent with the target sound. Specifically, and consistent with an attentional effect, the AtAmVt condition, the 3^rd^ PC amplitude evoked by timbre deviants presented in the attended coherent stream (AtVt) was significantly stronger than in the unattended incoherent stream (AmVt) (14-170 ms, cluster-level p_FWE_ < 0.001, T_max_ = 7.37). However, when the visual stimulus was coherent with the masker, the expected enhancement of the target stream was absent; there was no significant difference between the deviant-evoked 3^rd^ PC amplitude in the attended incoherent stream (AtVm) and in the unattended coherent stream (AmVm) (p_FWE_ > 0.05; Figure 6C).

**Figure 6.**
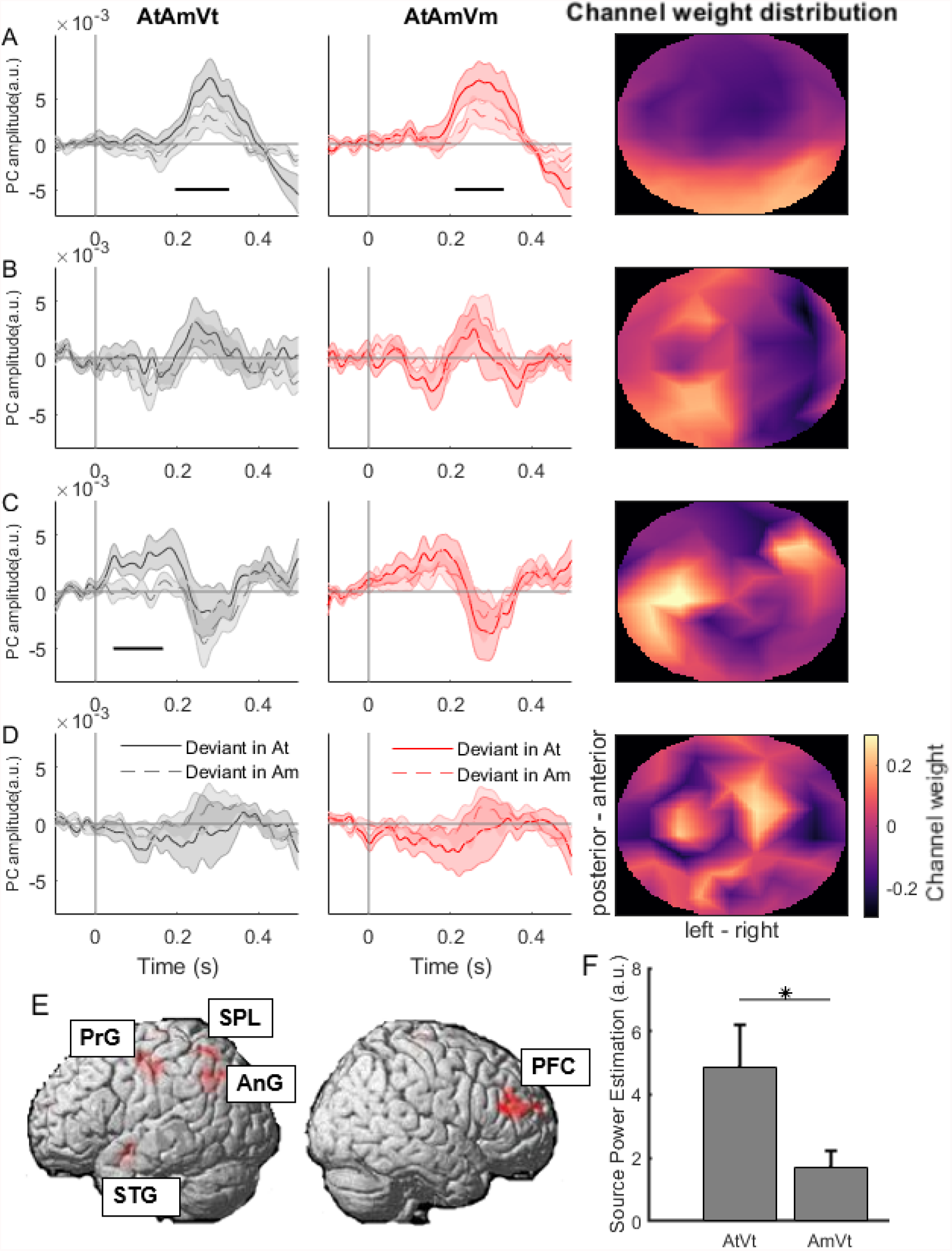
Attentional enhancement of deviant-evoked ERPs: principal component analysis. **(A**,**B**,**C**,**D)** Deviant-evoked response for the 1^st^ PC, 2^nd^ PC, 3^rd^ PC, and 4^th^ PC of ERP, respectively. The first two columns represent the visual coherence conditions AtAmVt, AtAmVm, respectively. Black bars indicate the time periods with a significant difference between At and Am. Shaded areas indicate SEM across subjects. The third column represents the spatial topography map of the principal component weights across channels. Color indicates the weight (warm: high, cool: low). **(E)** Source localization of the 3^rd^ PC (difference between deviants in At vs. Am embedded in the AtAmVt stream). **(F)** The mean source power estimates from the regions of interest for the AtVt and AmVt in the condition AtAmVt.

For the masker auditory stream, we also observed that, for PC3, the timbre deviants presented in the masker stream were modulated by audiovisual temporal coherence; deviants in AmVm evoked a stronger response (time lag between 100-132 ms, T_max_ = 3.79, cluster-level p_FWE_ < 0.001) than in AmVt. However, in the target auditory stream, the visual condition did not influence the magnitude of the deviant response; no significant difference was observed between the 3^rd^ PC amplitude evoked by deviants in the coherent stream (AtVt) and the incoherent stream (AtVm). The 3^rd^ PC was dominated by the responses from the left temporal and right frontal channels (Figure 6C, last column).

Source localization was carried out to identify the brain areas (Figure 6E) likely generating the difference in 3^rd^ PC response amplitude (difference between deviants in At vs. Am embedded in the AtAmVt stream; since the same difference in the AtAmVm stream was not significant, we did not subject it to source reconstruction). Overall, source localization of the 3^rd^ PC explained 88.13 ± 3.08% (mean ± SEM across participants) of sensor-level variance. We first identified the regions showing a significant response to deviants presented in the At (relative to the baseline). These regions included the left angular gyrus (AnG), superior parietal lobule (SPL), precentral gyrus (PrG), superior temporal gyrus (STG), and right/middle superior frontal gyrus (PFC) (see Table 1 for detailed results). We then compared source power estimates in this network between the two types of deviant responses. Pooling across all regions, the source power estimates significantly differed between the timbre deviants in the target and masker streams (Figure 5F, p < 0.001). When analyzed separately, all regions but one (STG, p = 0.06) showed significantly stronger source activity for AtVt deviants than AmVt deviants (all p < 0.05, uncorrected).

**Table 1.** Source localization results. Summary statistics of all clusters showing significant difference between At and Am in the 3^rd^ PC of the ERP, AtAmVt condition (omnibus F test: p_FWE_ < 0.05).

| Cluster label | Peak MNI coords | Voxel extent | T<sub>max</sub> | Z<sub>max</sub> | p values |
| --- | --- | --- | --- | --- | --- |
| Left angular gyrus (AnG) / superior parietal lobule (SPL) | -40 -58 40 / -32 -54 40 | 247 | 5.10 | 4.61 | 0.002 |
| Left precentral gyrus (PrG) | -34 -20 44 | 370 | 5.06 | 4.58 | 0.001 |
| Left superior temporal gyrus (STG) | -54 0 -10 | 49 | 4.64 | 4.25 | 0.016 |
| Left superior parietal lobule (SPL) | -32 -56 60 | 120 | 4.71 | 4.31 | 0.007 |
| Right superior frontal gyrus (prefrontal cortex, PFC) | 18 60 20 | 72 | 4.61 | 4.24 | 0.013 |
| Right middle frontal gyrus (PFC) | 42 44 18 | 367 | 5.04 | 4.56 | 0.001 |

### Correlations between behavioral performance and EEG

To examine the relationship between the EEG responses and behavioral performance, we calculated Pearson correlation coefficients between measures of behavioural performance and neural activity. Outliers were deleted using Cook’s distance if the distance was larger than 3 times the means of Cook’s distance. We first considered whether the magnitude of the deviant response in the target stream correlated with overall behavioural performance (mean d’ across all visual conditions), reasoning that participants with a stronger deviant response might be better able to accurately report timbre deviants. For both PC1 and PC3, the peak-to-peak PCs of ERP amplitudes obtained for the deviants in the target stream (At) correlated with overall d’ performance (PC1 peak-to-peak amplitude: Figure 7A, r = 0.55, p = 0.019; PC3: Figure 7B, r = 0.61, p = 0.005).

**Figure 7.**
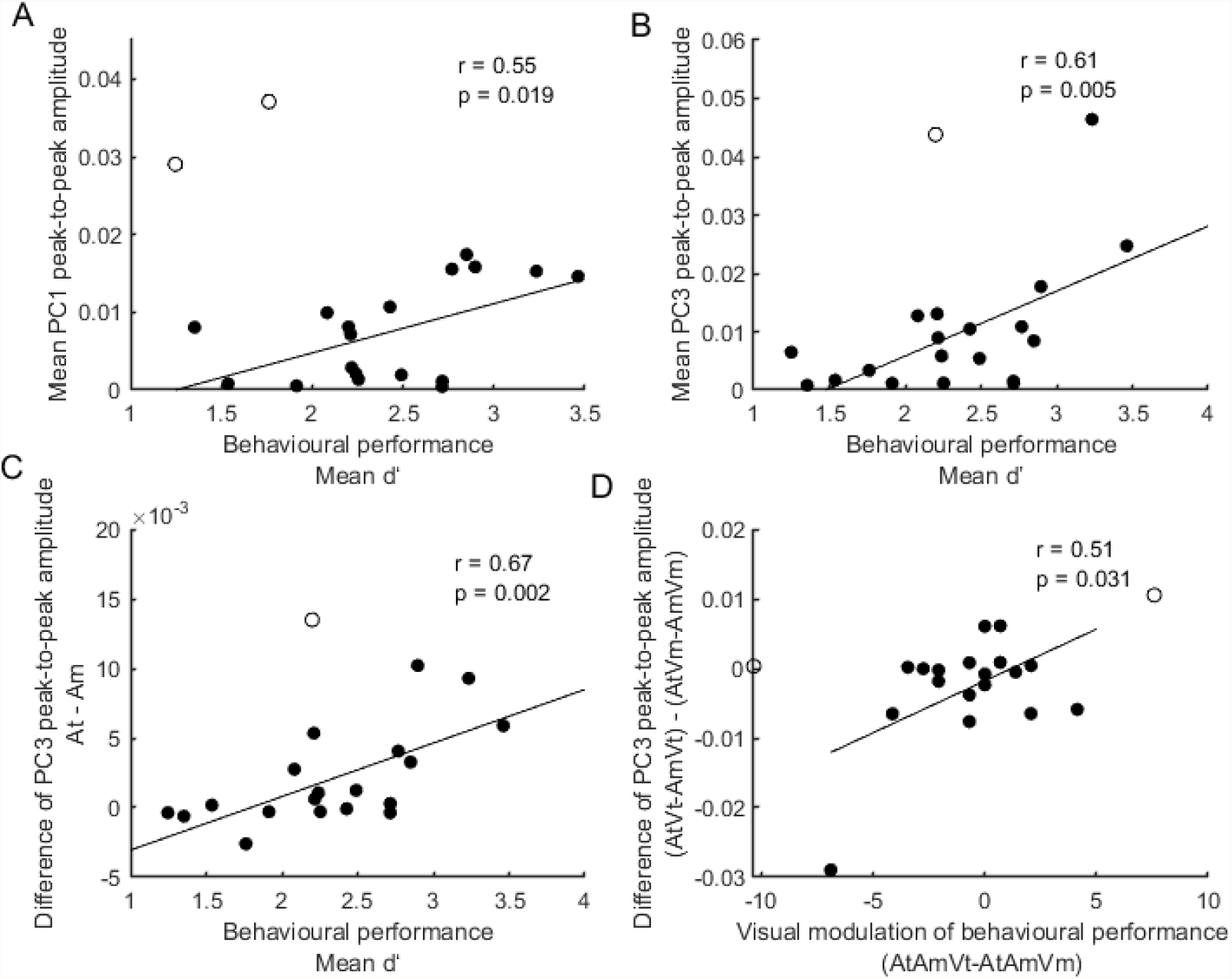
Correlations between the behavioral performance and EEG responses. **(A, B)** The correlation between mean d’ and the mean 1^st^ PC and 3^rd^ PC peak-to-peak amplitude over conditions (AtAmVt, AtAmVm, and AtAmVi), respectively. **(C)** The correlation between mean d’ and the mean 3^rd^ PC peak-to-peak amplitude (At - Am). **(D)** Visual coherence modulation of behaviour performance with EEG responses. The correlation between the hit rate difference (AtAmVt - AtAmVm) and the 3^rd^ PC peak-to-peak amplitude (AtVt – AmVt vs AtVm - AmVm). The unfilled circles represent outliers.

The auditory selective attention task required that participants not only detect timbre deviants, but that they successfully differentiated target and masker events. We therefore hypothesised that listeners who more successfully engaged selective attention mechanisms might show larger differences in the magnitude of deviant response to target and masker deviants. To test this we subtracted the peak to peak amplitude of EEG responses for masker deviants from the peak to peak amplitude to target deviants, and then measured the correlation between the EEG responses difference with the behavioral performance (d’). For PC1 there was no significant correlation (r=-0.01, p=0.971), but for PC3 there was (Figure 7C, r=0.67, p=0.001).

Finally, while the visual condition did not significantly influence behavioural performance at the group level, there was significant heterogeneity within our listeners. To determine whether modulation of behavioural performance by the visual stimulus correlated with the magnitude of the attention × visual condition interaction in PC3, we considered the difference in the normalised d’ performance for target-coherent and masker-coherent trials (i.e. the difference in target-coherent d’ and masker-coherent performance d’ / overall d’) and correlated this with the difference in the attentional modulation of the 3^rd^ PC peak-to-peak amplitude across visual conditions, i.e. AtVt-AmVt vs AtVm-AmVm (Figure 7D, r = 0.51, p = 0.031). While the correlation was significant and in the predicted direction (i.e. participants who showed a benefit for target-coherent trials had a greater attentional modulation in the target-coherent condition), we note that itss principally driven by a single participant whose removal renders the correlation non-significant.

## Discussion

This study used an auditory selective attention task, performed in the presence of a temporally modulated visual stimulus, to dissect the neural signatures of selective attention and audiovisual temporal coherence. Our EEG data of envelope responses reveal evidence for audiovisual integration of temporally coherent audiovisual envelopes which occurred independently of selective attention. In contrast, selective attention had a strong effect on the amplitude of TRFs derived from the envelope responses, with TRFs corresponding to target streams yielding higher amplitudes than those corresponding to masker streams. To further investigate audiovisual binding we examined the EEG responses to the timbre deviants which occurred independently of the amplitude envelopes of the audio(visual) streams. The fact that the EEG responses elicited by the timbre deviants were affected by the visual coherence of the stimulus can be interpreted as evidence that temporal coherence in the audiovisual streams favored the emergence of a fused audiovisual percept, which contrasts more strongly against the deviants than a purely auditory stream would. In direct support of this notion, we observed that, in some spatiotemporal components of the neural response, audiovisual temporal coherence interacted with selective attention.

### Temporal coherence facilitates AV integration independently of attention

Based on the stimulus envelope reconstruction analysis, we found that the cortical responses to the AV amplitude envelope were better explained by an AV integration model than by a linear summation (A+V) model. This was apparent in both the attended and unattended streams, suggesting attention was not required to link audio and visual streams. Our study thus provides evidence that temporal coherence between the auditory and visual stream facilitates AV integration independently of attention. This result is in contrast to previous studies using speech as stimuli. For example, Morís Fernández et al., (2015) studied fMRI responses to natural speech, and concluded that visual-auditory integration was only observed for attended speech streams. Using EEG, Ahmed et al., (2021) found broadly consistent results with the previous study, demonstrating that responses to attended speech were better explained by an AV model, while the responses to unattended speech were better explained by the A+V model. However, their integration model outperformed the linear summation model for unattended speech at very short (0-100ms lag) latencies, suggesting that distinct multisensory computations occur at different processing stages. In contrast to studies utilizing natural speech and videos of faces, our visual disc was much simpler. One possibility, which is already noted in Atilgan et al. (2018), is that bottom up audiovisual integration does occur independently of attention for simple non-speech stimuli. Another possibility is that watching a competing talker is more distracting than watching an uninformative disc, perhaps leading to observers actively suppressing a competing face in the context of a selective attention task. A final difference might be that subjects in Ahmed et al. (2021) were instructed to look at the eyes of the face, whereas our listeners fixated on the disc itself; potentially the radius changes of the disk, presented at the fovea, provide a more salient temporal cue. In support of this possibility, we note that the stimulus reconstruction accuracy of the visual-only decoder in the independent condition was quite high, and significantly larger than that of the audio-only decoder.

We used a forward model to examine the cortical representation of the sound amplitude envelope across all EEG channels. In the unattended sound stream, the TRF_AV_ amplitude was significantly stronger than the summation of TRF_A_ and TRF_V_ amplitude, which suggests that AV integration occurs pre-attentively. This result is consistent with our results from the envelope reconstruction (Figure 2), as well as a previous study from Crosse et al (2015), both in terms of the direction of the effect (AV vs. A+V) and its latency in the ∼200 ms range. In contrast, in the attended stream, the linear summation model and the integration model performed equivalently. Furthermore, attention strongly modulated the TRF, with the TRF_AV_ amplitude for the target stream being significantly larger than that for the masker stream. This finding is consistent with previous studies, demonstrating an enhancement of attended speech streams (Ding & Simon, 2012; Mesgarani & Chang, 2012) and audiovisual streams (Zion Golumbic et al., 2013). An open question is why audiovisual temporal coherence did not influence the attended stream TRF_AV._ Perhaps the enhancement of the TRF by attention generated a ceiling effect, or possibly if we had required subjects to attend to the visual stimulus we might have observed stronger audiovisual interactions. Nevertheless, our TRF results suggest the temporal coherence facilitates AV integration, and attention further enhances the TRF amplitude of coherent AV streams.

### Attention and coherence effects on the deviant evoked responses

In this study, we adapted the behavioral paradigm of previous studies (Atilgan & Bizley, 2021; Maddox et al., 2015) to examine the effects of attention and coherence on the neural responses to the change of an independent auditory feature (timbre here). However, we failed to replicate the behavioural findings of Maddox et al. (2015) and Atilgan and Bizley (2021). Two key differences may explain this: first, in adapting the task to a wider participant sample (including Cantonese or English listeners), the magnitude of the timbre deviants was increased, as pilot studies suggested this would be necessary. However, in the event, this effectively rendered the task easier, and the overall d’ scores are higher in the current dataset than in previous ones. The second difference was that, in these previous studies, listeners were also required to detect occasional colour deviants in the visual stimulus, which required them to maintain some level of attention towards the visual modality. In our version of the experiment, the visual stimulus neither contained deviants of its own, nor did it provide cues that might facilitate the detection of auditory deviants. It is possible that this difference explains why, at the group level, we did not observe a significant effect of audiovisual temporal coherence on auditory deviant detection.

A whole-scalp analysis of deviant-evoked ERPs brought evidence for a main effect of attention. Attended deviants evoked stronger responses than unattended deviants, independent of whether they were presented in a stream that was coherent or incoherent with visual stimulation. A principal component analysis based on the timbre deviant elicited ERPs revealed interactions between attention and audiovisual temporal coherence. The 3^rd^ PC showed an attention dependent enhancement of the responses embedded in the target stream, only when the visual stimulus was coherent with the target stream. When the visual stimulus was coherent with the masker stream, responses evoked by timbre deviants were similar in both target and masker streams (Figure 6C). The neural responses to a non-binding feature (timbre) in the unattended stream was mediated by the coherent feature (envelope), providing evidence that temporal coherence facilitates binding to create an AV object. PC3 was also correlated with differences in behavioural task performance, with the magnitude of attentional modulation scaling with overall behavioural performance d’ (Figure 7C). There was some evidence for a correlation between the extent to which the visual condition influenced behavioural performance and the magnitude of the temporal coherence dependent attentional effects (Figure 7D), although this requires replication, preferably in the context of task parameters that more reliably elicit a modulation of task performance by audiovisual temporal coherence.

Our results are consistent with previous studies on ‘cocktail party effect’ speech stream segregation, in which congruent visual stimuli enhanced the cortical representation of the speech envelope of attended speech streams relative to unattended streams (Crosse, Di Liberto, & Lalor, 2016; Golumbic et al., 2013). However, unlike in these previous studies, where visual speech provided temporal and contextual information about the auditory envelope, in our study we used a simple disc as a visual stimulus, which provided no information about the auditory deviant. While previous studies have demonstrated that attention dedicated to one feature of an object may enhance the responses to other features of the object in both auditory (Alain & Arnott, 2000; Maddox & Shinn-Cunningham, 2012; Shamma et al., 2011; Shinn-Cunningham, 2008) and visual modalities (Blaser et al., 2000; O’Craven et al., 1999), our results provide new evidence that temporal coherence modulates the attentional enhancement of the neural response to the timbre deviants (“other” features) of the AV object.

We used source reconstruction to localize the brain areas showing interaction between attention and temporal coherence, and the brain areas included the superior temporal cortex and several regions in the frontoparietal network. The superior temporal sulcus (Beauchamp et al., 2004; Calvert & Campbell, 2003; Kayser & Logothetis, 2009) and the superior parietal lobule (Molholm et al., 2006) have previously been shown to be involved in AV object formation. Similarly, the superior frontal cortex receives inputs from multimodal areas involved in the identification of objects (M Petrides & Pandya, 1999; Michael Petrides & Pandya, 2002), while the medial prefrontal cortex (PFC) receives auditory and visual information from the superior temporal sulcus and superior temporal cortex (Kondo et al., 2003, 2005; Michel & Morales, 2020). Thus, our source localization results are consistent with the notion that the synergistic interaction between attention and coherence is mediated by areas downstream from sensory regions, primarily in the frontoparietal network.

In summary, in this study we examined the temporal coherence and attention effect on neural responses to the continuous sound envelope and the deviant evoked response, respectively. Temporal coherence facilitated the audiovisual integration pre-attentively, and attention further enhanced the audiovisual integration of the coherent audiovisual stream. Attention amplified a large portion of the deviant evoked response independent of temporal coherence, however, we also identified a spatiotemporal component of the responses, likely originating from the superior temporal gyrus and the frontoparietal network, in which both attention and coherence synergistically modulated deviant evoked responses. These results provide evidence for partly dissociable neural signatures of bottom-up (coherence) and top-down (attention) effects in AV object formation.

## Materials and Methods

### Participants

Twenty volunteers were recruited for this experiment (median ± standard deviation (SD) age, 22 ± 2 years; 12 males; 19 right-handed). All participants were healthy, had self-reported normal hearing and normal or corrected-to-normal vision. Prior to the experiment, each participant gave written informed consent. All procedures were approved by the Human Subjects Ethics Sub-Committee of the City University of Hong Kong.

### Stimuli

We adapted the behavioral paradigm from previous psychoacoustics studies (Atilgan & Bizley, 2021; Maddox et al., 2015). Stimuli included two simultaneously presented auditory streams and one visual stream. One auditory stream was meant to be attended, and will be referred to as the target sound (At), while the other one was meant to be unattended, and will be referred to as the masker stream (Am). Finally, stimulation included a concurrently presented visual stream (V) which comprised a radius-modulated disc. Auditory streams were independently amplitude-modulated and the modulation of the visual disc could be temporally coherent either with the amplitude of the target stream (AtAmVt), the masker stream (AtAmVm), or independent of both (AtAmVi) (Figure 1A).

The envelopes below 7 Hz were generated using the same methods as in Maddox et al. (2015). Each auditory stream consisted of one continuous amplitude modulated synthetic vowel, either /u/ or /a/, which were generated by filtering a click train at four “formant” frequencies (F1-F4). The fundamental frequency (F0) of vowel /u/ was 175 Hz, and the formant peaks were 460, 1105, 2975, 4263 Hz, while the F0 of vowel /a/ was 195 Hz, and the formant peaks were 936, 1551, 2975, 4263 Hz. Auditory deviants were embedded in the auditory streams by temporarily changing the timbre of the vowel. The deviant in vowel /u/ transitioned (in F1/F2 space) towards the vowel /ε/, with the maximum timbre change resulting in formant peaks at 525, 1334, 2975, 4263 Hz, while the deviant in vowel /a/ transitioned towards /i/ with formant peaks at 860, 1725, 2975, 4263 Hz. The duration of each deviant was 200 ms, which included a linear change of the formants towards the deviant for 100 ms and then back for 100 ms.

The visual stimulus was a grey disc surrounded by a white ring presented at the center of the black screen. The radius of the visual stimulus was modulated by the visual envelope, such that the disc subtended between 1° and 2.5°, and the white ring extended 0.125° beyond the grey disc.

Each trial lasted 14 s. The target audio stream and the visual stream were each 14 s in duration. The masker stream began 1s later and lasted 13 s. The initial 1 s, during which only the target stream was audible, provided the cue for the listener which was the to-be-attended target stream. Auditory deviants could occur at any time during a window beginning 2 s after the onset of the target audio stream and ending 1 s before the end of the trial, subject to the constraint that the minimum interval between auditory deviants was 1.2 s. On average each stream contained 2 deviants (range 1-3 across trials). Unlike Maddox et al. (2015), the visual stream did not contain any colour deviants.

### Procedure

Participants were seated in a sound-attenuated room. Auditory stimuli were presented binaurally via earphones (ER-3, Etymotic Research, Elk Grove Village, IL, USA), using an RZ6 signal processor at a sampling rate of 24414 Hz (Tucker-Davis Technologies, Alachua, FL, USA). The sound level was calibrated at 65 dB SPL. Visual stimuli were presented on a 24-inch computer monitor using the Psychophysics Toolbox for MATLAB. Participants were asked to pay attention to the target auditory stream and to detect the embedded auditory deviants by pressing a keyboard button. They were instructed to refrain from pressing buttons in response to any events in the masker stream.

Before the actual task, all participants completed a training session to verify that they were able to detect the auditory deviants. The training session included four blocks, and each block included 9 trials. The feedback of performance was given after each block, and all participants showed they could perform the experiment (d’ > 0.8) in at least one block of four.

Participants were instructed to minimize eye blinks and body movements during the EEG recording. Continuous EEG signals were collected using an ANT Neuro EEGo Sports amplifier from 64 scalp channels at a sampling rate of 1024 Hz. The EEG signals were grounded at the nasion and referenced to the Cpz electrode. Each participant completed 12 blocks of the task, with 18 trials (6 trials x 3 conditions) in each block. Trials belonging to different conditions were presented in a randomly interleaved order. In total, each participant completed 216 trials (72 trials x 3 conditions). Feedback on behavioral performance was provided after each block. Triggers corresponding to trial and deviant onset were recorded along with the EEG signal.

### Behavioral data analysis

A ‘hit’ was defined as the response to the deviant in the target auditory stream within 1 s following the onset of the deviant, and a ‘false alarm’ was defined as the response to a deviant that occurred in the masker stream. To study how visual coherence affects auditory deviant detection, we conducted a one-way repeated measures ANOVA on the sensitivity measure d’ with a within-subjects factor of AV condition (visual coherent with the target, AtAmVt, visual coherent with the masker, AtAmVm, and independent visual AtAmVi).

### EEG signal pre-processing

EEG signals were pre-processed using the SPM12 Toolbox (Wellcome Trust Centre for Neuroimaging, University College London) for MATLAB. Continuous data were downsampled to 500 Hz, high-pass filtered at a cut-off frequency of 0.01 Hz, notch-filtered between 48 Hz and 52 Hz, and then low-pass filtered at 90 Hz. All filters were fifth-order zero-phase Butterworth. Eyeblink artifacts were detected using the signal in the channel Fpz, and removed by subtracting the first two spatiotemporal components associated with each eyeblink from all channels (Nicole et al., 2002). The EEG data were then re-referenced to the average of all channels. The preprocessed data were further analyzed in two ways: For the response to the sound amplitude envelope, the pre-processed data were bandpass filtered between 0.3 and 30 Hz (Crosse et al., 2015), downsampled to 64 Hz, and subjected to a calculation of the TRF, or used for stimulus reconstruction (see below). For the deviant evoked response analysis, the pre-processed data were epoched from -100 ms to 500 ms relative to deviant onset. Epoched EEG signals were then denoised using the “Dynamic Separation of Sources” (DSS) algorithm (de Cheveigné & Simon, 2008), which is commonly used to maximize reproducibility of stimulus-evoked response across trials and maintain the differences across the different stimulus types (here: 2 vowel types × 3 experimental conditions). Epoched data were linearly detrended, and the first seven DSS components were preserved and applied to project the data back into sensor space. Denoised data were averaged across trials.

### EEG response to sound amplitude envelopes

#### Stimulus reconstruction

To investigate how visual temporal coherence and attention affect multisensory integration, we quantified the accuracy of neural tracking of the sound amplitude envelope. We reconstructed amplitude envelopes of different elements of the AV scene (Crosse et al., 2015) based on the EEG data using a linear model as follows:

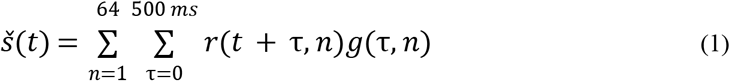

where *Š* (*t*) is the reconstructed envelope; *r*(*t* + *τ,n*)is the EEG data at channel *n*; and *g* is the linear decoder representing the linear mapping from the response to stimulus amplitude envelope at time lag *τ*. The time lag *τ* ranged from 0 to 500 ms post-stimulus. The decoder was obtained separately for each condition using ridge regression as follows:

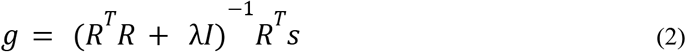

where *R* is the lagged time series of the EEG response, *λ* is the ridge parameter, *I* is the regularization term, and *s* is the sound amplitude envelope. The decoder is a multivariate impulse response function computed from all channels and all time-lags simultaneously. Decoders corresponding to the AV, A-only, and V-only streams were generated separately as follows:

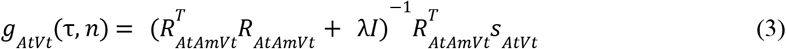

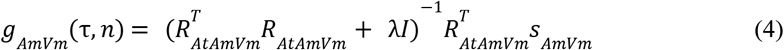

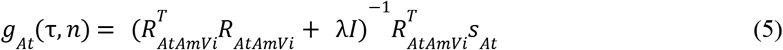

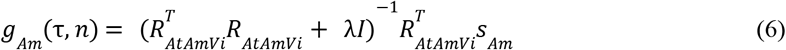

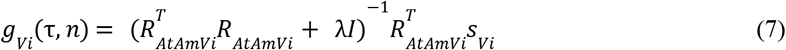

Since in the condition AtAmVi, the envelope of At, Am, and Vi are independent of each other, we could obtain the decoder of the envelopes of the auditory target only, auditory masker only, and visual only, respectively. To obtain the decoder for each condition, we used leave-one-out cross-validation to select the *λ* value (from the set of 10^−6^, 10^−5^,…, 10^5^, 10^6^) for which the correlation between *Š* (*t*)and *s(t)* is maximized. To assess the effect of AV integration, we reconstructed the sound envelopes (both target and masker sound) using the integration AV decoder and the algebraic sum of the A and V decoder (A+V), separately, based on the following formulas:

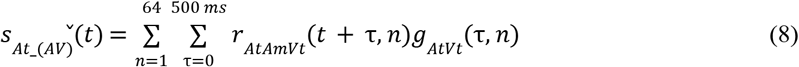

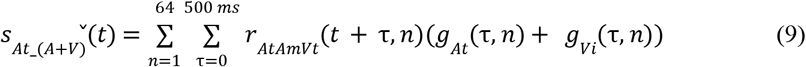

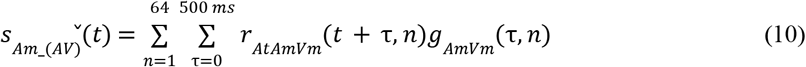

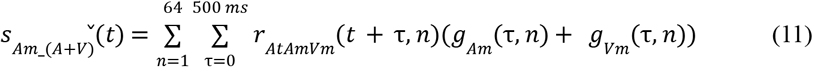

The reconstruction accuracy (r) was defined as the Pearson correlation coefficient between the actual stimulus envelope and the estimated envelope.

To test whether the reconstruction accuracy using either the AV decoder (“integration model”) and/or A+V decoder (“summation model”) was significantly larger than chance, we conducted a nonparametric permutation test. The null distribution of 1000 Pearson’s r values was created for each subject by calculating the correlation between randomly shuffled response trials of estimated sound envelopes and actual sound envelopes. We estimated sound envelopes using each decoder separately, and generated the null distribution for each condition. Comparisons across conditions (target vs. masker) and across decoders (AV vs. A+V) were performed using Wilcoxon signed-rank tests, and multiple comparisons were corrected using Bonferroni correction.

### Single-lag analysis

The decoder *g*(τ,*n*)in the above analysis was run on the EEG signal over a 500 ms convolution lag (i.e., information from all lags were used to construct the decoder). In a follow-up analysis, to quantify the contribution of individual time lag towards reconstruction, we trained the decoders on EEG at single lags from 0 to 500 ms (O’Sullivan et al., 2015). Since the sampling rate of the response was 64 Hz (33 points in the 0-500 ms latency range), we generated 33 separate decoders. The reconstruction formula at each time lag becomes:

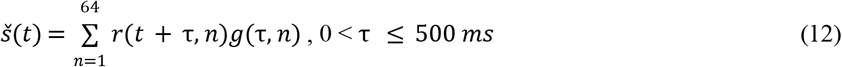

We reconstructed the sound amplitude envelope (target and masker) using the single-lag AV decoder and A+V decoder separately. To compare the single-lag reconstruction (AV vs A+V) both for the target and masker conditions, we converted the single-lag reconstruction accuracy into one-dimensional images (time) using the SPM toolbox. The converted images were then subjected to statistical inference using paired t-tests in SPM12. The significance threshold was set at p <0.05 (two-tailed), and p values were corrected using a false-discovery-rate (FDR) approach at an FDR = 0.05.

### Temporal response function (TRF) estimation

To investigate how the visual temporal coherence and attention affect AV integration across the EEG channels, we estimated the linear temporal response function (TRF) (Crosse, Di Liberto, Bednar, et al., 2016) which links the EEG response at each channel and the sound envelope. The TRF is the linear filter that best describes the brain’s transformation of the sound envelope to the continuous neural response at each EEG channel location (Haufe et al., 2014). TRFs were estimated separately for each experimental condition (AtAmVt, AtAmVm, AtAmVi) as follows:

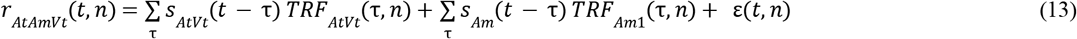

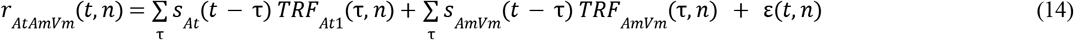

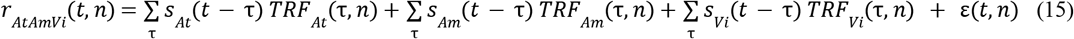

Where *r*_*AtAmVt*_, *r*_*AtAmVm*_, and *r* _*AtAmVi*_ are the EEG response in each of the 3 conditions respectively; *t* is time, *n* is the index of the EEG channel under consideration; *s* _*At*_, *s*_*Am*_, and *s*_*Vi*_ are the stimulus envelopes of At, Am, and Vi, respectively; *τ* represents the convolution time lag (−100 ms to 500 ms), and *ε*(*t,n*)is the residual “error”, that is, the part of the EEG recording not explained by the TRF model. We use the term *TRF*_*At*1_ to describe the TRF in the AtAmVt condition and *TRF*_*Am*1_ in the AtAmVm condition to differentiate them from the *TRF*_*At*_ and *TRF*_*Am*_ estimated from the AtAmVi condition. The TRF for each condition was calculated at time lags from -100 ms to 500 ms relative to the stimulus as follows:

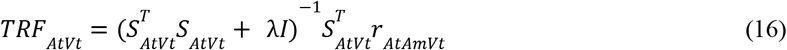

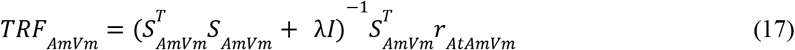

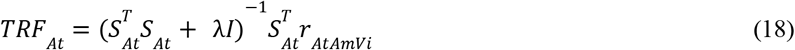

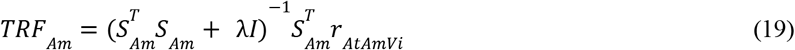

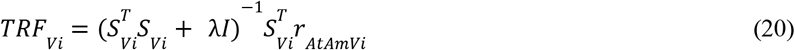

where ***S*** is the lagged time series of the stimulus envelope; *I* is the regularization term used to prevent overfitting; and *λ* is the ridge parameter. *TRF*_*AtVt*_, *TRF*_*AmVm*_, *TRF*_*At*_, *TRF*_*Am*_, and *TRF*_*Vm*_ were fitted separately for each condition using the MATLAB toolbox adapted from a previous study by Crosse et al. (2016). The TRF of each channel was estimated using leave-one-out cross-validation. The best *λ* (in the range of 2^10^, 2^11^,…, 2^21^) was selected based on the maximum correlation coefficient between the predicted response with the actual neural response for each channel. The EEG signal of each trial (13 s long) was used to estimate the TRF, modeling the neural response to the simultaneous presentation of both At and Am.

To test whether the AV integration occurs in target and/or masker streams, we compared the TRF amplitude between the temporally coherent and independent conditions for any EEG channel and time point. Single-participant TRF data were converted into three-dimensional images (2D: spatial topography, 1D: time) and entered into paired t-tests between two coherence conditions (coherent: AV; independent: A+V), separately for two attentional conditions (AtVt, target; AmVm, masker). In the analysis of the AtAmVt condition (visual stream coherent with the target sound), the coherent AV stream corresponds to *TRF*_*AtVt*_. This TRF is compared against the sum of two TRFs (*TRF*_*At*_ + *TRF*_*Vi*_) in the condition AtAmVi, in which the auditory stream and visual stream are independent: one based on the same sound (At) presented in a temporally independent context (*TRF*_*At*_), and one based on the visual stream presented in a temporally independent context (*TRF*_*Vi*_). Similarly, in the analysis of the AtAmVm condition (visual stream coherent with the masker sound), the coherent stream corresponds to *TRF*_*AmVm*_, which is compared against *TRF*_*Am*_ + *TRF*_*Vi*_ estimated from the independent condition AtAmVi. We also compared the TRF amplitude between the two temporally coherent conditions (*TRF*_*AtVt*_ vs *TRF*_*AmVm*_) to examine how attention affects AV integration.

The three paired t-tests were implemented in SPM12. The resulting spatiotemporal maps of t-statistics were thresholded at p < 0.05 (two-tailed), and then corrected for multiple comparisons across spatiotemporal voxels at a family-wise error (FWE)-corrected p = 0.05 (cluster-level) under random field theory assumptions (Kilner et al., 2005).

### Auditory Deviant-evoked ERP

To assess how attention and visual coherence affect deviant-evoked activity, the EEG data were first subject to a traditional channel-by-channel mass-univariate analysis. Epoched data were averaged over trials, separately for the deviants in At and Am and for each visual condition (Vt, Vm, Vi). Single-subject ERP data were converted into three-dimensional images (two spatial dimensions and one temporal dimension) and entered into a repeated-measures ANOVA with two within-subjects factors: attention (attended: deviant in the At stream, unattended: deviant in the Am stream) and visual coherence (coherent with the sound containing deviants: deviants in AtVt and AmVm; visual condition independent of the sound: AtVm and AmVt). The two-way repeated-measures ANOVA was implemented as a GLM in SPM12. The resulting statistical parametric maps, representing the main and interaction effects, were thresholded at p < 0.05 (two-tailed) and corrected for multiple comparisons across spatiotemporal voxels at a family-wise error (FWE)-corrected p = 0.05 (cluster-level).

In a follow-up attempt to isolate dissociable neural signatures of attention and visual coherence, we concatenated the ERP data across participants and used principal component analysis (PCA) to reduce the EEG data dimensionality and obtain spatial principal components (PCs, representing the weight of channel topographies) and temporal principal components (representing voltage time-series). The first four PCs (explaining 80% of the original variance across participants) were used to extract single-participant ERP components for further analysis. Each PC was then converted into one-dimensional images (time) and subject to statistical inference using repeated-measures ANOVAs, as above. Significance thresholds were kept identical to the traditional univariate analysis, but correction for multiple comparisons was implemented across time points (rather than spatiotemporal voxels).

To further infer the most probable cortical sources contributing to the 3^rd^ PC, for which we have identified significant main and interaction effects of attention and coherence (see Results), we projected the PC time series back into sensor space (using the corresponding spatial PC as a weight matrix) as a basis for subsequent source reconstruction. To this end, we used the multiple-sparse-priors method of source localization under group constraints (Litvak & Friston, 2008) as implemented in SPM12. Single-subject source estimates corresponding to the At and Am responses (50-150 ms after the deviant onset) and the corresponding baseline (−100 to 0 ms relative to deviant onset) were converted into 3D images in Montreal Neurological Institute (MNI) space, smoothed with a 6 × 6 × 6 mm Gaussian kernel, and entered into a repeated-measures ANOVA with two within-subjects factors (condition: deviants evoked responses in coherent target AtVt vs. deviants evoked responses in incoherent masker AmVt; time window: evoked response vs. baseline). Only two conditions were included in the source localization since we only observed a significant ERP difference between these two conditions (see Results). To restrict the number of regions, we performed a region of interest analysis by first identifying those regions showing a significant difference between the evoked response and the baseline for the deviants in the AtVt condition, and then compared the source estimates between the responses evoked by the deviants in the AtVt and AmVt in those regions. First, the statistical parametric map of the main effect of stimulus vs. baseline, corresponding to a difference between At and Am source estimates during the evoked response and their respective baselines, was thresholded at a family-wise error (FWE)-corrected p = 0.05 (cluster-level). The Neuromorphometrics atlas implemented in SPM12 was used to label the sources. Then, within this network (pooling across all regions as well as for each region separately), we compared the source activity estimates between At and Am deviants using paired t-tests.

### Correlating timbre deviant evoked ERP magnitude with behavioral performance

Since the behavioral task was to detect deviants in the target auditory stream, we extracted the EEG responses to deviants in At and measured the peak to peak amplitude of the PCs of ERP identified above. We then calculated the Pearson correlation coefficients between the behavioral mean d’ and the mean PC amplitude over conditions (AtAmVt, AtAmVm, and AtAmVi). For the 1^st^ PC for which we have identified the significant main effect of attention (see Results), the negative and positive peaks were measured between 100 to 200 ms, and 220 to 300 ms. respectively. For the 3^rd^ PC for which we have identified significant main and interaction effects of attention and coherence (see Results), the positive and negative peaks were measured between 50 to 160 ms, and 220 to 400 ms, respectively. Prior to calculating the correlations, we fitted the behavioral performance d’ with the PC peak-to-peak amplitude using a linear regression model, and detected the outliers in each condition using Cook’s distance (threshold: 3 means of Cook’s distance).

## Acknowledgements

We thank Ruben Reuben Chaudhuri and On-mongkol Jaesiri for assistance with data collection. JKB was funded by a Wellcome / Royal Society Sir Henry Dale Fellowship (098418/Z/12/Z). This work has been supported by the European Commission’s Marie Skłodowska-Curie Global Fellowship (750459 to R.A.) and a grant from European Community/Hong Kong Research Grants Council Joint Research Scheme (9051402 to R.A. and J.S.).

